## Supplementary File for "Intratumoral delivery of FLT3L with CXCR3/CCR5 ligands promotes XCR1^+^ cDC1 infiltration and activates anti-tumor immunity"

Supplementary information includes:

Supplementary Figures 1-10

Supplementary Table 1 – Resource table.

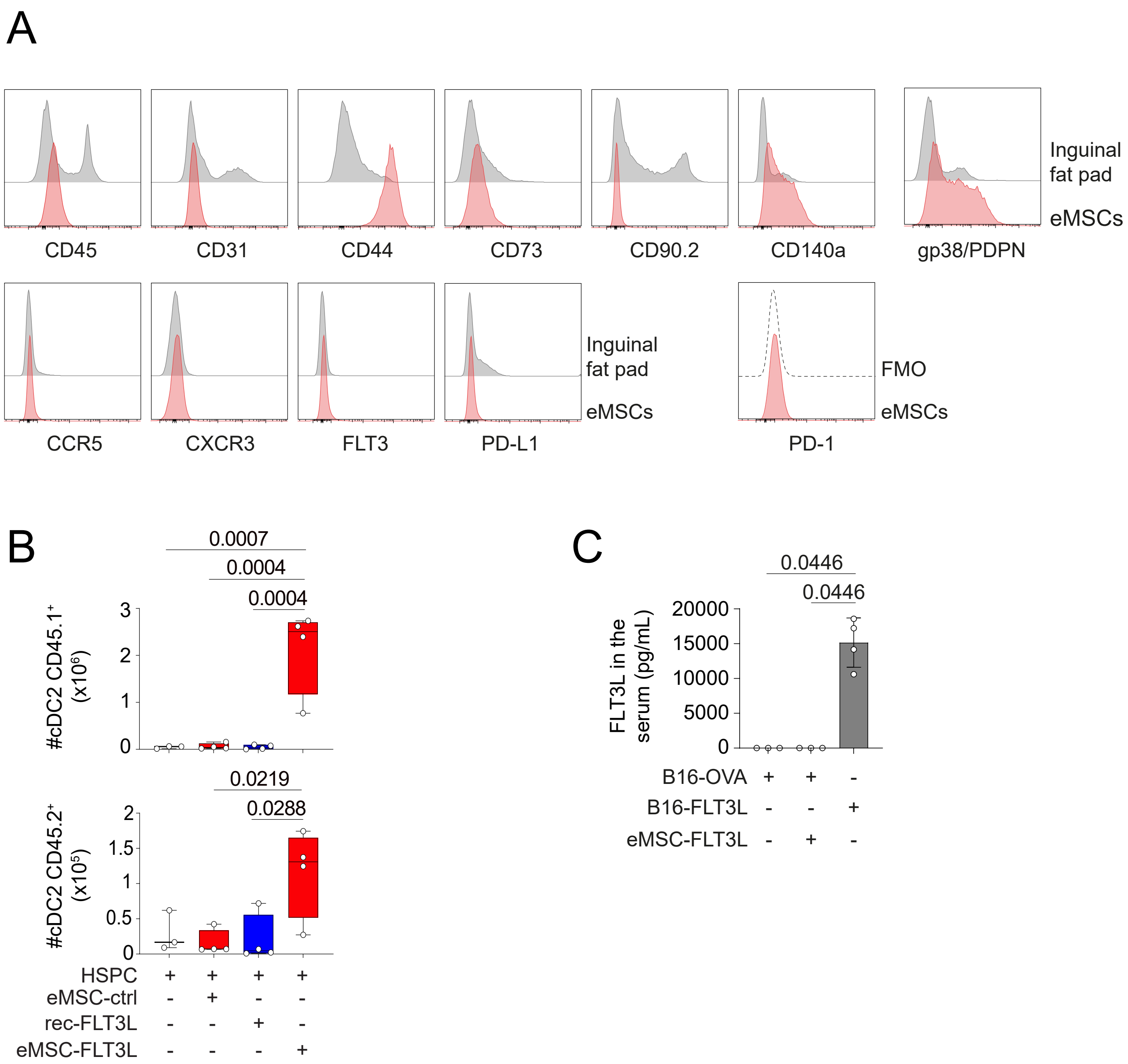

**<u>Supplementary Figure 1</u>: Phenotypic characterization and biological activity of eMSCs**

**(A)** Histograms showing flow cytometry phenotypic analysis of eMSCs or inguinal fat pad cells using various classical and conventional markers for stromal and stem cells.

**(B)** Quantification at day 12 of CD45.1 and CD45.2 cDC2s in the synthetic niches containing HSPC only (n=3 plugs) or with eMSC-FLT3L, eMSC-Ctrl or recombinant FLT3L (rec-FLT3L) (n=4 plugs), one experiment, one-way ANOVA-test with Tukey's multiple comparisons. Box plots show median, 25th–75th percentiles, minimum–maximum whiskers, with all data points displayed.

**(C)** Circulating FLT3L levels measured by ELISA in the serum of mice bearing B16-OVA (n=3), B16-OVA+eMSC-FLT3L (n=3) or B16-FLT3L (n=4), 4 days after eMSCs injection, one experiment, Kruskal-Wallis test with Dunn's multiple comparisons.

**(B-C)** Source data are provided as a Source Data file.

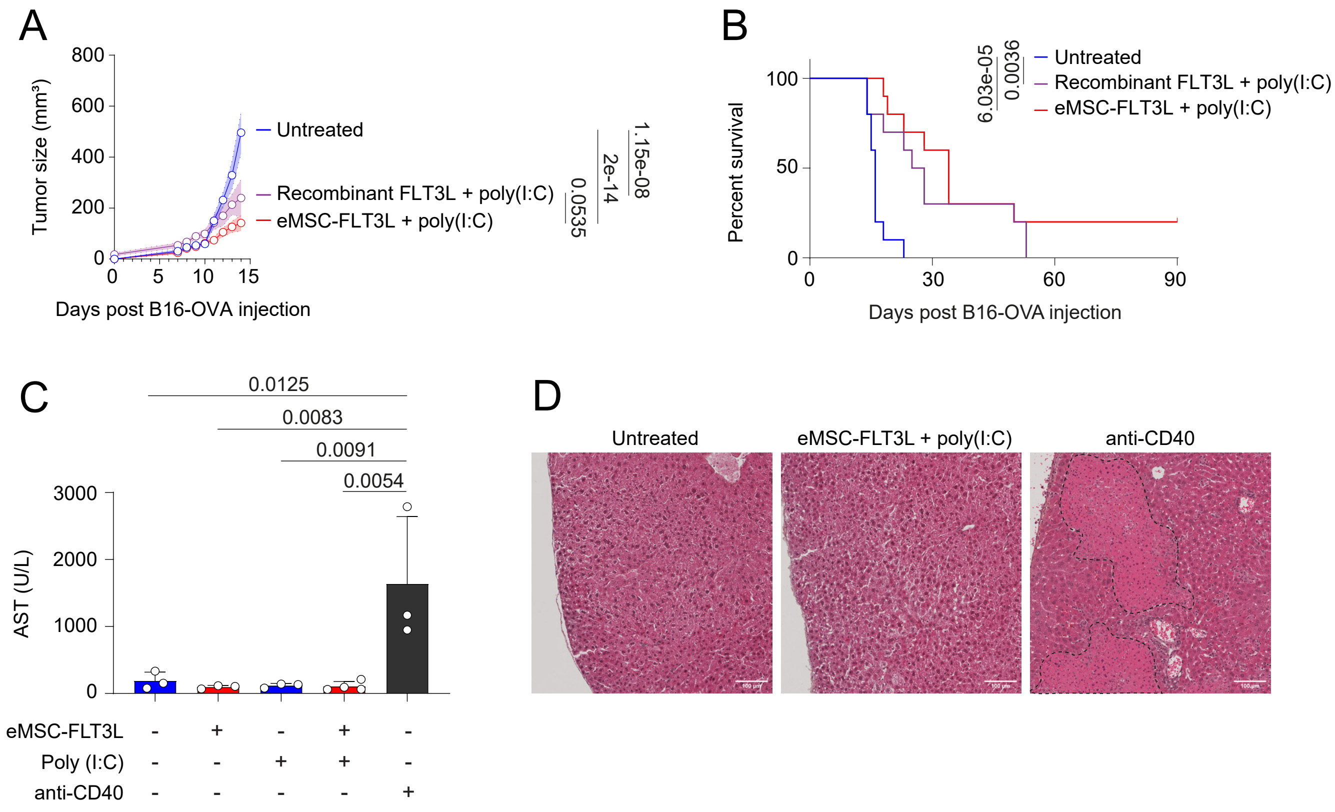

<u>**Supplementary Figure 2:**</u> **eMSC-FLT3L + poly(I:C) immunotherapy is more effective than recombinant FLT3L + poly(I:C) treatment, without inducing toxicity.**

**(A)** Tumor growth curves until day 14, n=10 mice per group, two independent experiments, two-way ANOVA-test with Tukey's multiple comparisons test. A line represents the mean and SEM is shown with the colored area.

**(B)** Survival curves, n=10 mice per group, two independent experiments, log-rank (Mantel-Cox) test.

(**C**) Level of AST in serum collected 48h after poly(I:C) or anti-CD40 treatment, n=4 (eMSC-FLT3L+poly(I:C)), n=3 mice (other ones), one experiment, one-way ANOVA-test with Tukey's multiple comparisons test.

(**D**) H&E staining of fixed liver tissue from mice treated with eMSC-FLT3L + poly(I:C) therapy or anti-CD40. Necrotic lesions are shown with the dashed black lines.

**(A-C)** Source data are provided as a Source Data file.

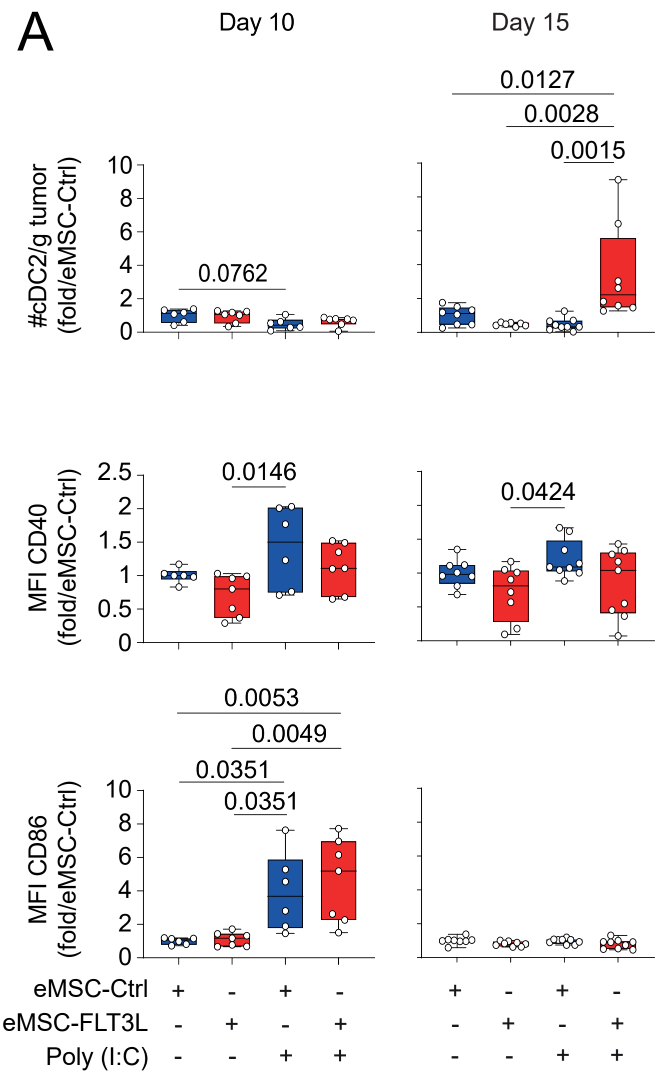

**<u>Supplementary Figure 3:</u> Intratumoral delivery of eMSC-FLT3L + poly(I:C) stimulates cDC2 infiltration in tumor.**

**(A)** Quantification of the absolute number of intratumoral cDC2/g tumor and the mean fluorescence intensity (MFI) of CD40 and CD86. Results are shown as fold change to control (eMSC-Ctrl). Day 10: n=6 (eMSC-Ctrl, eMSC-Ctrl+poly(I:C)), n=7 (eMSC-FLT3L, eMSC-FLT3L+poly(I:C)) mice per group, two independent experiments, one-way ANOVA-test with Tukey's multiple comparisons. Day 15: n=8 (eMSC-Ctrl, eMSC-FLT3L+poly(I:C)), n=7 (eMSC-FLT3L), n=9 (eMSC-Ctrl+poly(I:C) mice per group for the number of cDC2s/g tumor; n=8 (eMSC-Ctrl, eMSC-FLT3L), n=9 (eMSC-Ctrl+poly(I:C), eMSC-FLT3L+poly(I:C)) mice per group for the MFI, two independent experiments, one-way ANOVA-test with Tukey's multiple comparisons. Box plots show median, 25th–75th percentiles, minimum–maximum whiskers, with all data points displayed. Source data are provided as a Source Data file.

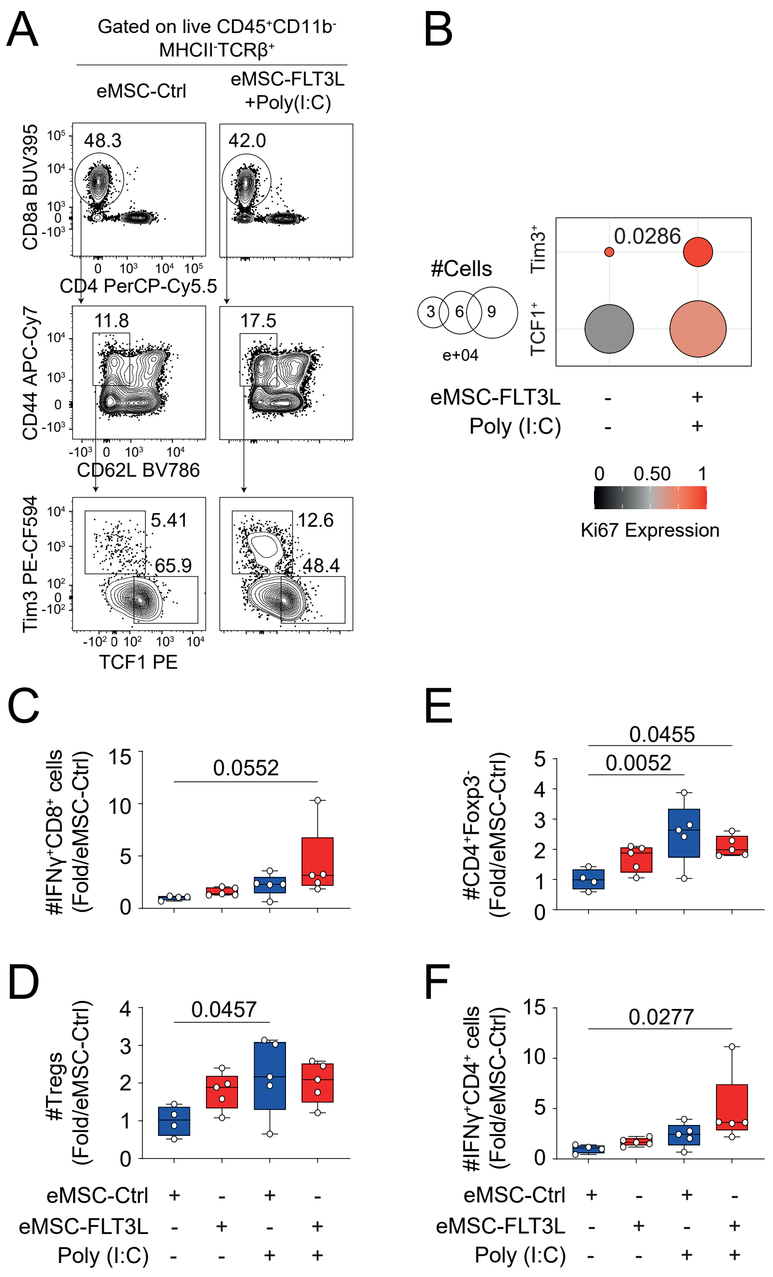

**Supplementary Figure 4: Intratumoral delivery of eMSC-FLT3L + poly(I:C) increase proliferative stem-like CD8$^+$ T cells, IFNγ$^+$CD8$^+$ and IFNγ$^+$CD4$^+$ but not T regulatory cells.**

**(A)** Representative flow cytometry plots of CD8 T cells in the tumor-draining lymph node at day 15.

**(B)** Quantification of the absolute number and Ki67 expression of exhausted (Tim3$^+$) and stem-like (TCF1$^+$) T cells. Dots represent the absolute number of cells while the colors represent the expression level of Ki67. n=4 mice per group, one experiment, statistics are done on the absolute numbers, two-tailed Mann-Whitney test.

**(C-F)** Quantification of the absolute number of IFNγ$^+$CD8$^+$ T cells (C), CD4$^+$Foxp3$^+$ T regulatory cells (Tregs) (D), CD4$^+$Foxp3$^-$ cells (E) and IFNγ$^+$CD4$^+$ cells (F) at day 15. n=4 (eMSC-Ctrl), n=5 (other groups) mice per group, one experiment, one-way ANOVA-test with Dunnett's multiple comparison test. Box plots show median, 25th–75th percentiles, minimum–maximum whiskers, with all data points displayed.

**(B-F)** Source data are provided as a Source Data file.

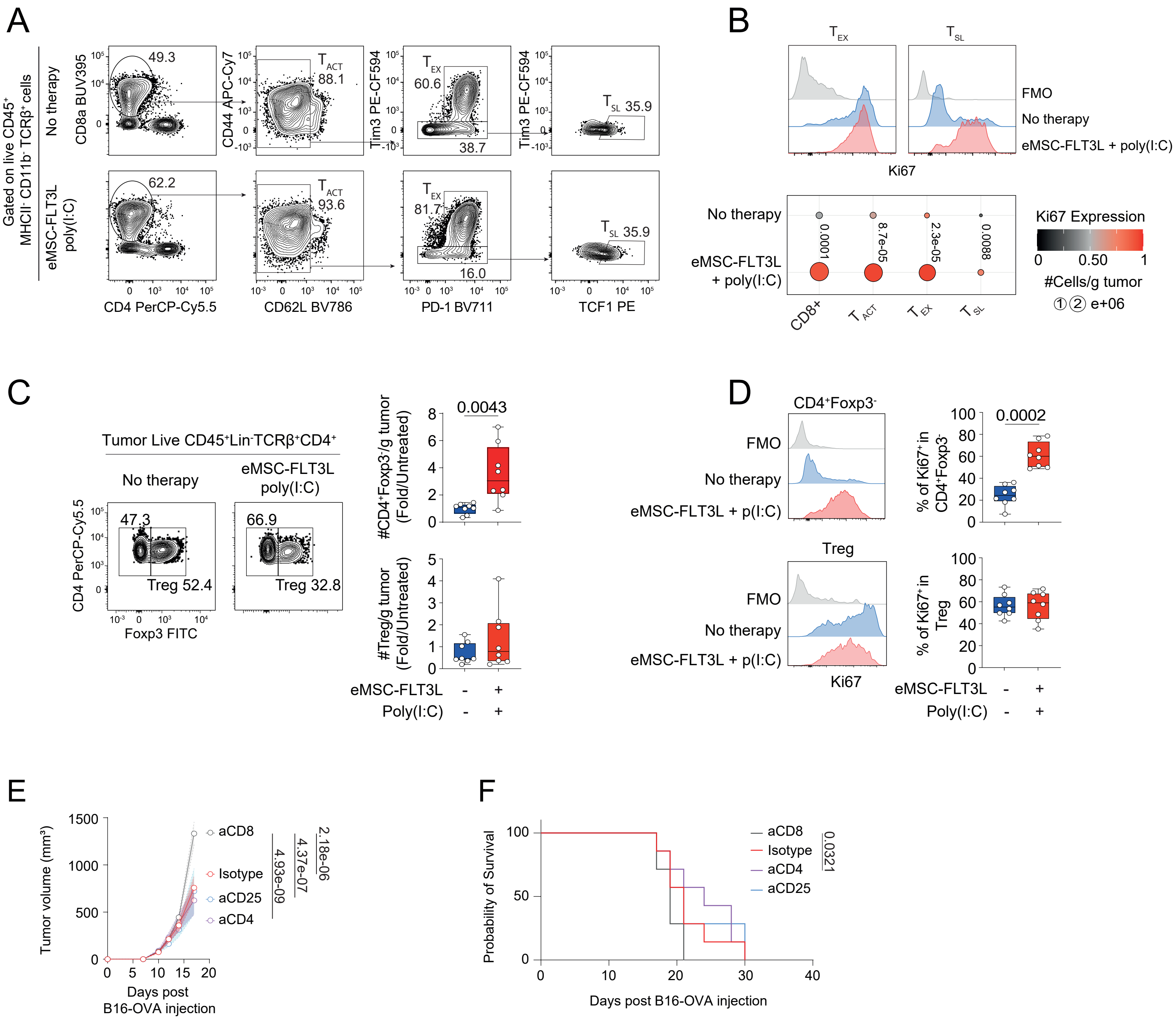
A
B
C
D
E
F
Gated on live CD45+ MHCII- CD11b- TCRβ+ cells
No therapy
eMSC-FLT3L poly(I:C)
CD8a BUV395
CD4 PerCP-Cy5.5
CD44 APC-Cy7
CD62L BV786
Tim3 PE-CF594
PD-1 BV711
TCF1 PE
49.3
62.2
TACT 88.1
TACT 93.6
TEX 60.6
TEX 81.7
38.7
16.0
TSL 35.9
TEX
TSL
FMO
No therapy
eMSC-FLT3L + poly(I:C)
Ki67
0.0001
8.7e-05
2.3e-05
0.0088
CD8+
TACT
TEX
TSL
Ki67 Expression
#Cells/g tumor
e+06
Tumor Live CD45+Lin-TCRβ+CD4+
CD4 PerCP-Cy5.5
Foxp3 FITC
47.3
Treg 52.4
66.9
Treg 32.8
#CD4+Foxp3-/g tumor (Fold/Untreated)
0.0043
#Treg/g tumor (Fold/Untreated)
eMSC-FLT3L
Poly(I:C)
CD4+Foxp3-
Treg
eMSC-FLT3L + p(I:C)
% of Ki67+ in CD4+Foxp3-
0.0002
% of Ki67+ in Treg
Tumor volume (mm3)
Days post B16-OVA injection
aCD8
Isotype
aCD25
aCD4
2.18e-06
4.37e-07
4.93e-09
Probability of Survival
0.0321

**Supplementary Figure 5: Intratumoral delivery of eMSC-FLT3L + poly(I:C) stimulates infiltration and proliferation of CD4+ T cells but not of T regulatory cells.**

**(A-B)** Representative flow cytometry plots (A) and quantification (B) of activated ($T_{ACT}$), exhausted ($T_{EX}$), and stem-like ($T_{SL}$) $CD8^+$ T cells. Dots represent the absolute number of cells/g tumor while the colors represent the expression level of Ki67. n=4 mice per group, one experiment, statistics are done on the absolute numbers, two-tailed unpaired t-test.

**(C)** Representative flow cytometry plots and absolute number quantification of $CD4^+Foxp3^-$ and Treg. Results are shown as fold change to control (Untreated)**.** n=8 mice per group, two independent experiments, two-tailed unpaired t-test.

**(D)** Representative flow cytometry plots and frequency of $Ki67^+$ cells in $CD4^+Foxp3^-$ cells and Treg**.** n=8 mice per group, two independent experiments, two-tailed unpaired t-test.

**(C-D)** Box plots show median, 25th–75th percentiles, minimum–maximum whiskers, with all data points displayed.

**(E)** Tumor growth curves. n=7 mice per group, one experiment, two-way ANOVA-test with Tukey's multiple comparisons. A line represents the mean and SEM is shown with the colored area.

**(F)** Survival curves. n=7 mice per group, one experiment, log-rank (Mantel-Cox) test.

**(B-F)** Source data are provided as a Source Data file.

A

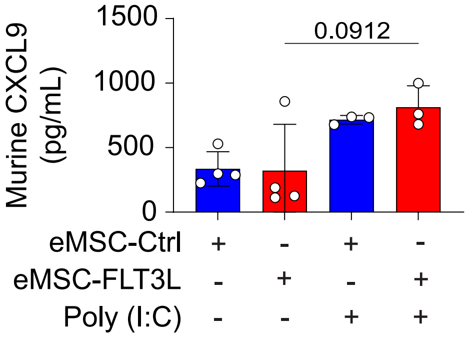

**Supplementary Figure 6: poly(I:C) monotherapy is sufficient to upregulate CXCL9 expression in tumor homogenates.**

(**A**) Murine CXCL9 ELISA done on tumor homogenates 24h after poly(I:C) injection. n=3 (eMSC-Ctrl+poly(I:C), eMSC-FLT3L+poly(I:C)), n=4 (eMSC-Ctrl, eMSC-FLT3L), one experiment, Kruskal-Wallis test with Dunn's multiple comparisons. Data are presented as mean values ± SD. Source data are provided as a Source Data file.

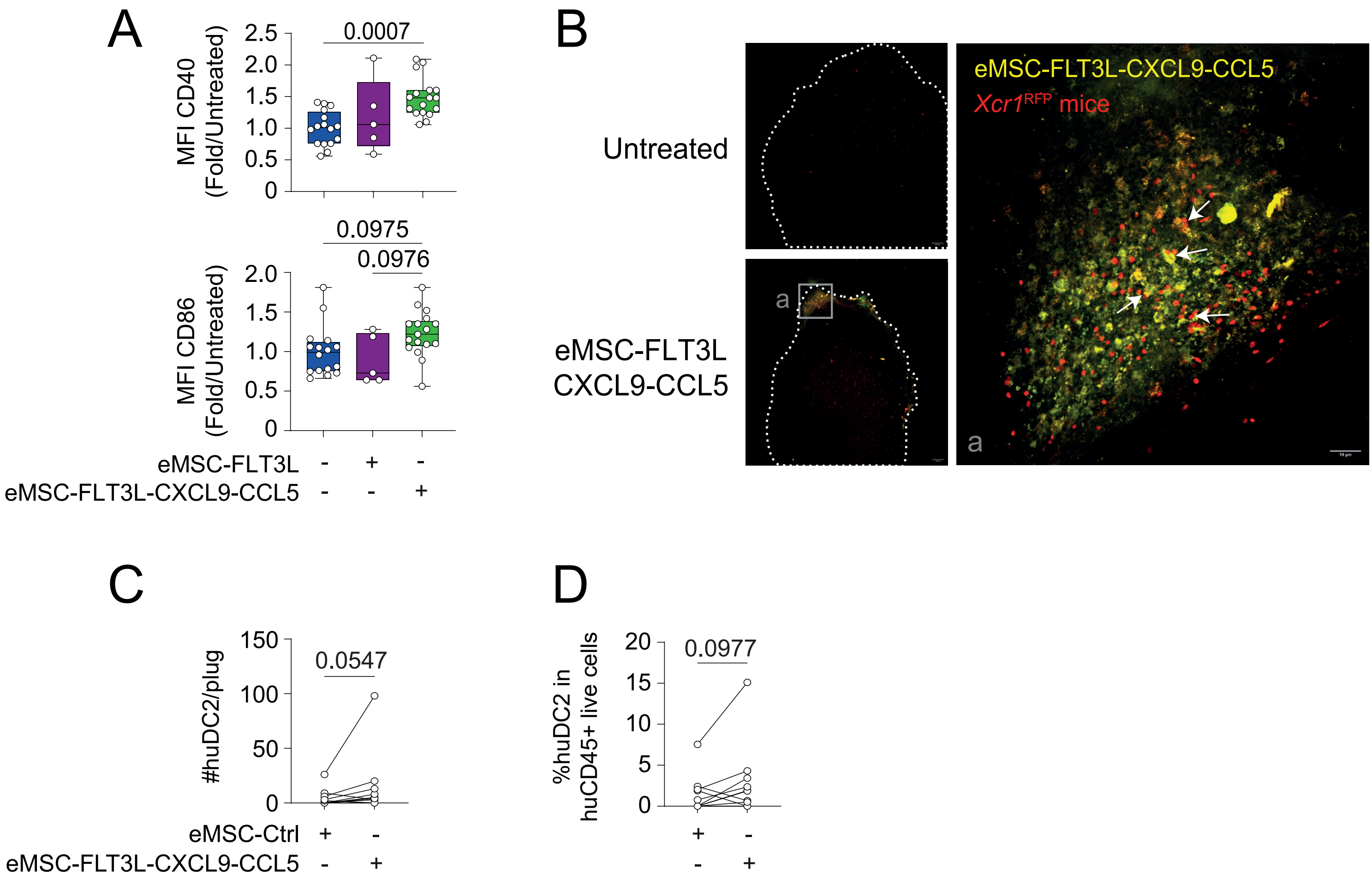

**Supplementary Figure 7: eMSC-FLT3L-CXCL9-CCL5 are localized at the border of the tumor and attract hucDC2s in synthetic niches in reconstituted BRGSF mice.**

**(A)** Quantification by flow cytometry of the mean fluorescence intensity (MFI) of CD40 and CD86 at day 17. Results are shown as fold change to control (Untreated). n=16 (untreated), n=17 (eMSC-FLT3L-CXCL9-CCL5), n=5 (eMSC-FLT3L) mice per group, three independent experiments, one-way ANOVA with Tukey's multiple comparisons test. Box plots show median, 25th–75th percentiles, minimum–maximum whiskers, with all data points displayed.

(**B**) Immunofluorescence imaging of YUMM-OVA tumors injected in *Xcr1*$^{RFP}$ mice. These tumors received 3 injections of DMEM (Untreated) or eMSC-FLT3L-CXCL9-CCL5. Images were taken with a spinning disk 24h after the last injection. cDC1s are in red, while eMSC-FLT3L-CXCL9-CCL5 are in yellow. Tumors are delimited with the doted white lines.

(**C-D**) Quantification of the absolute number (C) and frequencies (D) of human cDC2/plug. n=16 plugs per group, three independent experiments, two-tailed Wilcoxon matched-pairs signed rank test.

**(A, C-D)** Source data are provided as a Source Data file.

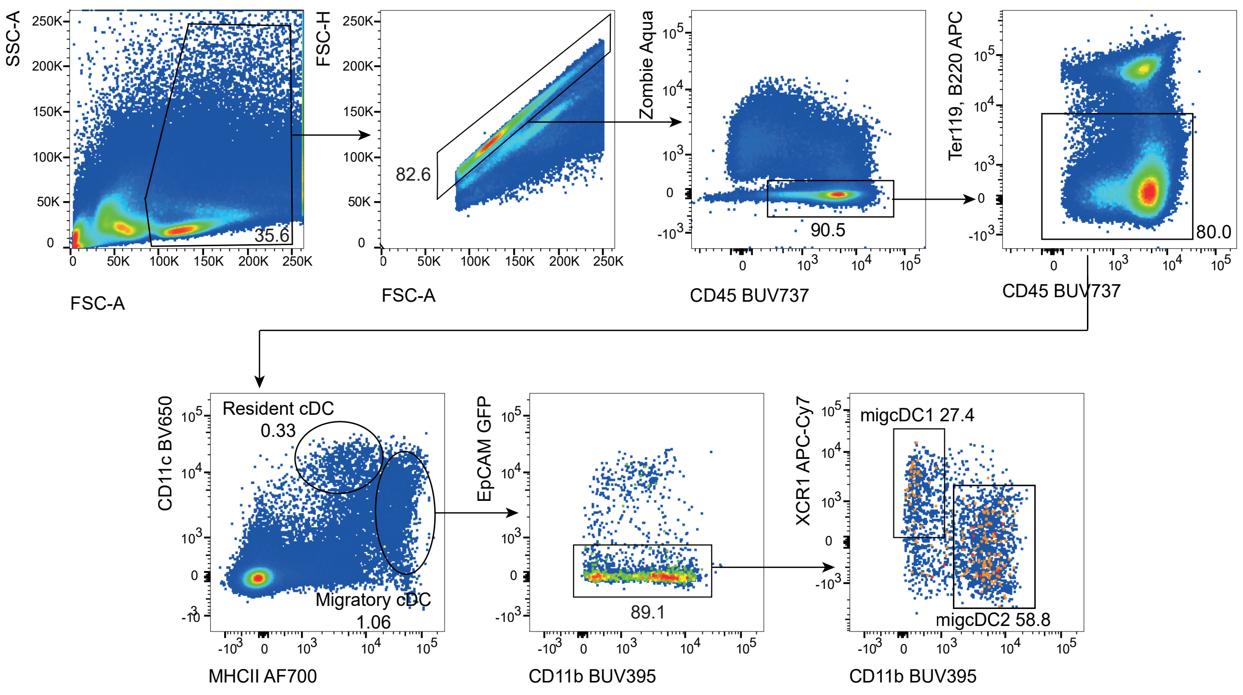

**Supplementary Figure 8: Gating strategy used to identify migratory cDC1s and cDC2s within tumor draining lymph nodes.**

**(A)** Gating strategy used to identify migratory cDC1s and cDC2s within tumor draining lymph nodes in Figure 1L and 4C.

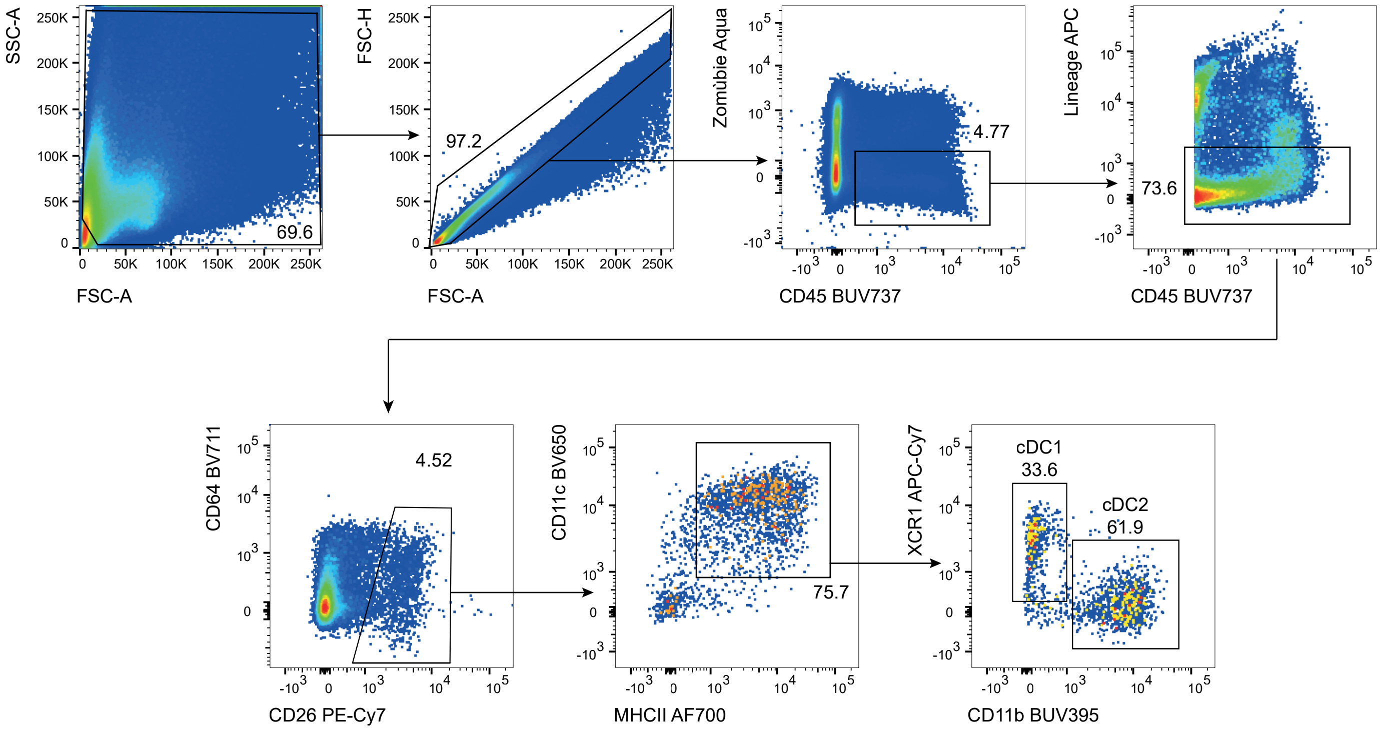

**Supplementary Figure 9: Gating strategy used to identify cDC1s and cDC2s within tumors.**

**(A)** Gating strategy used to identify cDC1s and cDC2s within tumors in Figure 1K, 3B-C, 6B-C, 6I, 7G-H and 7L. Lineage: Ter119, CD3, NK1.1, Ly6G, SiglecF, B220.

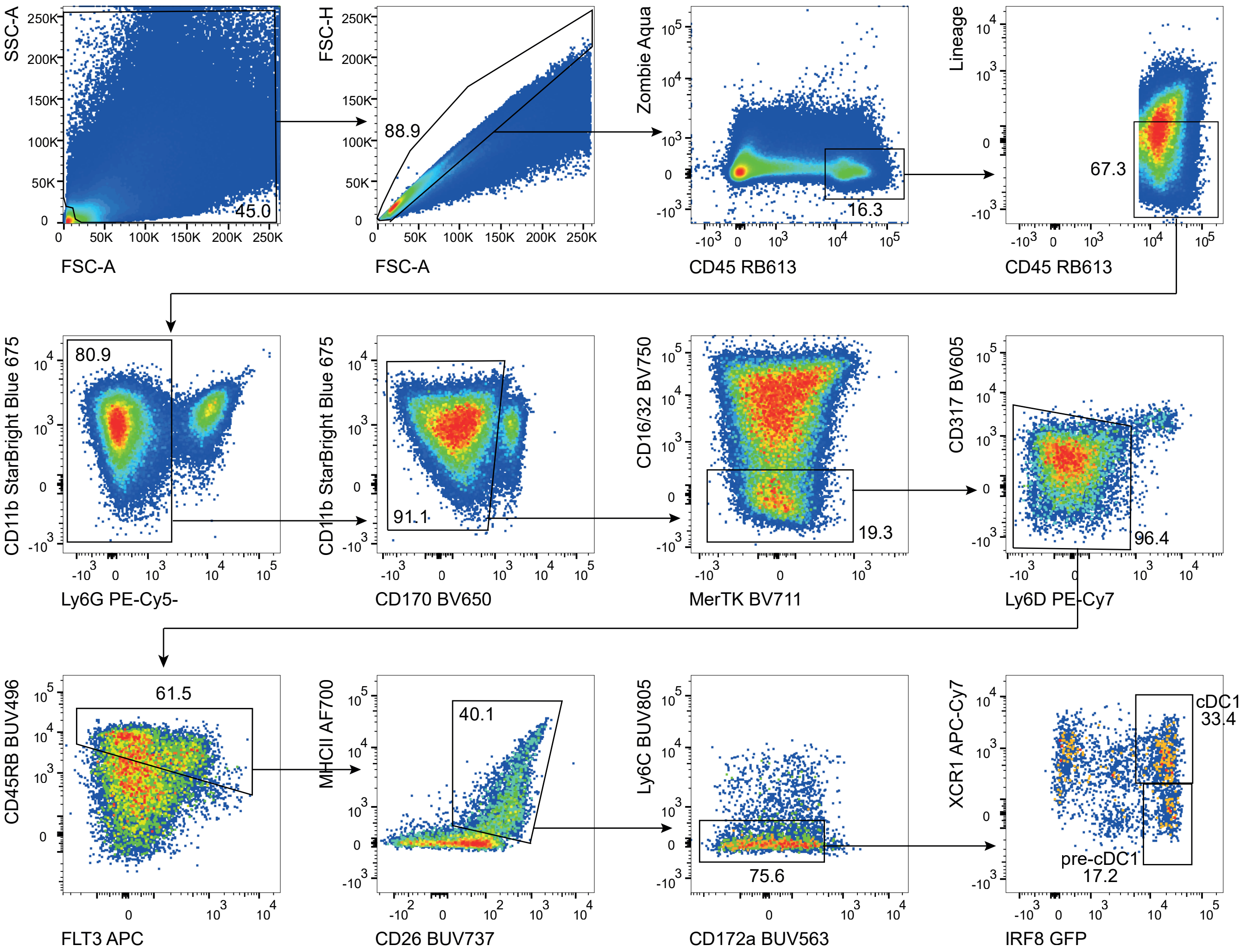

**Supplementary Figure 10: Gating strategy used to identify pre-cDC1s within tumors.**

**(A)** Gating strategy used to identify pre-cDC1s within tumors in Figure 7E. Lineage: Ter119, CD3, NK1.1, CD19.

**Supplementary Table 1:** **Resource table**.

| REAGENT or RESOURCE | SOURCE | IDENTIFIER | DILUTION |
|---|---|---|---|
| **Mouse antibodies** | | | |
| CD11b AF700 (M1/70) | Biolegend | Cat# 101222, RRID:AB_493705 | 200 |
| CD11b BV421 (M1/70) | Biolegend | Cat# 101236, RRID:AB_11203704 | 200 |
| CD11b FITC (M1/70) | Biolegend | Cat# 101206, RRID:AB_312789 | 200 |
| CD11b BUV395 (M1/70) | BD | Cat# 565976, RRID:AB_2721166 | 200 |
| CD11b BV785 (M1/70) | Biolegend | Cat# 101243, RRID:AB_2561373 | 200 |
| CD11b StarBright Blue 675 (5C.6) | Bio-Rad | Cat#MCA711SBB675, RRID:AB_3101464 | 200 |
| CD11c PeCy7 (N418) | Biolegend | Cat# 117318, RRID:AB_493568 | 200 |
| CD11c PE/Dazzle™ 594 (N418) | Biolegend | Cat# 117348, RRID:AB_2563655 | 200 |
| CD127 BV421 (A7R34) | Biolegend | Cat# 135024, RRID:AB_11218800 | 400 |
| CD16/32 BV510 (93) | Biolegend | Cat# 101333, RRID:AB_2563692 | 200 |
| CD24 BV510 (M1/69) | BioLegend | Cat# 101831, RRID: AB_2563894 | 400 |
| CD26 PeCy7 (H194-112) | Biolegend | Cat# 137810, RRID:AB_2564312 | 200 |
| CD3e PE (145-2c11) | Biolegend | Cat# 100308, RRID:AB_312673 | 100 |
| CD4 BV510 (RM4-5) | Biolegend | Cat# 100559, RRID:AB_2562608 | 400 |
| CD4 PercpCy5.5 (RM4-5) | Biolegend | Cat# 100540, RRID:AB_893326 | 400 |
| CD4 AF700 (RM4-5) | Biolegend | Cat# 100536, RRID:AB_493701 | 400 |
| CD44 APC/Fire750 (IM7) | Biolegend | Cat# 103062, RRID:AB_2616727 | 200 |

| | | | |
|---|---|---|---|
| CD45 BUV395 (30-F11) | eBioscience | Cat# 363-0451-82, RRID:AB_2925264 | 800 |
| CD45 BUV737 (30-F11) | eBioscience | Cat# 367-0451-82, RRID: AB_2895963 | 800 |
| CD45 APC/Fire750 (30F11) | Biolegend | Cat# 103154, RRID:AB_2572116 | 800 |
| CD45 RB613 (I3/2) | BD | Cat # 758191, RRID:AB_3690340 | 400 |
| CD45 AF700 (30-F11) | Biolegend | Car# 103128, RRID:AB_493715 | 800 |
| CD45 BUV805 (30F11) | BD | Cat# 568336, RRID:AB_3684191 | 800 |
| CD45.1 PeCy7 (A20) | Biolegend | Cat# 110729, RRID:AB_1134170 | 600 |
| CD45.1 BV605 (A20) | Biolegend | Cat# 110738, RRID:AB_2562565 | 600 |
| CD45.2 PerCp-Cy5 (104) | eBioscience | Cat# 45-0454-82 RRID: AB_953590 | 200 |
| CD45RB APC (C363-16A) | BioLegend | Cat# 103319; RRID: AB_2565228 | 200 |
| CD49a AF647 (Ha31/8) | BD | Cat# 562113, RRID:AB_11153312 | 200 |
| CD62L FITC (MEL-14) | Biolegend | Cat# 104406, RRID:AB_313093 | 200 |
| CD62L BV785 (MEL-14) | Biolegend | Cat# 104440, RRID:AB_2629685 | 200 |
| CD64 BV711 (X54-5/7.1) | Biolegend | Cat# 139311, RRID:AB_2563846 | 200 |
| CD80 BV421 (16-10A1) | Biolegend | Cat# 104725, RRID:AB_10900989 | 400 |
| CD8a (KT15) FITC | BioRad | Cat# MCA609G, RRID: AB_321407 | 400 |
| CD8a BV785 (53-6.7) | Biolegend | Cat# 100750, RRID:AB_2562610 | 400 |
| CD8b BUV395 (H35-17.2) | BD | Cat# 740278, RRID:AB_2740017 | 400 |
| CX3CR1 BV785 (SA011F11) | Biolegend | Cat# 149029, RRID:AB_2565938 | 200 |

| | | | |
|---|---|---|---|
| Ki67 APC (SolA15) | Invitrogen | Cat# 56-5698-82, RRID:AB_2637480 | 200 |
| KLRG1 PeCy7 (2F1) | Biolegend | Cat# 138416, RRID:AB_2561736 | 400 |
| Lag3 PE/Dazzle™ 594 (C9B7W) | Biolegend | Cat# 125224, RRID:AB_2572082 | 200 |
| Ly6C APC/ Fire750 (HK1.4) | Biolegend | Cat# 128046, RRID:AB_2616731 | 200 |
| Ly6C BUV805 (HK1.4) | BD | Cat# 755202, RRID:AB_3687609 | 200 |
| Ly6G BV650 (1A8) | Biolegend | Cat# 127641, RRID:AB_2565881 | 400 |
| I-A/I-E APC (M5/114.15.2) | Biolegend | Cat# 107614, RRID:AB_313329 | 400 |
| I-A/I-E AF700 (M5/114.15.2) | Biolegend | Cat# 107622, RRID:AB_493727 | 400 |
| PD-1 (CD279) BV711 (29F.1A12) | Biolegend | Cat# 135231, RRID:AB_2566158 | 200 |
| SiglecH BUV395 | BD | Cat# 747669, RRID: AB_2744230 | 200 |
| SiglecF BV421 (E50-2440) | BD | Cat# 562681, RRID:AB_2722581 | 400 |
| TCRb BV650 (H57-597) | Biolegend | Cat# 109251, RRID:AB_2810348 | 400 |
| Tim3 PE (RMT3-23) | BD | Cat# 568428; RRID: AB_345377 | 200 |
| PD-L1 BV605 | Biolegend | Cat# 124321, RRID:AB_2563635 | 400 |
| XCR1 PE (ZET) | Biolegend | Cat# 148204, RRID:AB_2563843 | 200 |
| XCR1 BV785 (ZET) | Biolegend | Cat# 148225, RRID:AB_2783119 | 200 |
| NK1.1 PE-Cy7 (clone S17016D) | Biolegend | Cat# 156513; RRID:AB_2888852 | 200 |
| CD40 PE-Dazzle584 (3/23) | Biolegend | Cat# 124629; RRID: AB_2572184 | 200 |
| CD40 RY610 (3/23) | BD | Cat# 759052, RRID:AB_3691150 | 200 |
| CD86 FITC (GL-1) | Biolegend | Cat# 105005, RRID: AB_313148 | 200 |
| CD86 PE (GL-1) | Biolegend | Cat# 105008, RRID:AB_313151 | 200 |
| CD86 AF700 (GL-1) | Biolegend | Cat#105023, RRID:AB_493720 | 200 |

| | | | |
|---|---|---|---|
| NKp46 BV421 (29A1.4) | Biolegend | Cat# 137612, RRID:AB_2563104 | 200 |
| EpCAM BV510 (G8.8) | Biolegend | Cat# 101831; RRID: AB_2563894 | 400 |
| Podoplani/gp38 BV421 (8.1.1) | Biolegend | Cat#127423 ; RRID: AB_2814017 | 200 |
| CD31 BV785 (390) | Biolegend | Cat#102435 ; RRID: AB_2810334 | 400 |
| ICAM-1 PE (YN1/1.7.4) | Biolegend | Cat#116107 ; RRID: AB_313698 | 400 |
| IFNy BV785 (XMG1) | Biolegend | Cat# 505837; RRID: AB_2629667 | 100 |
| IFNy BV650 (XMG1) | Biolegend | Cat# 505831; RRID: AB_11142685 | 100 |
| TCF1/TCF7 PE (S33-966) | BD | Cat# 564217; RRID: AB_2687845 | 100 |
| Foxp3 FITC (FJK-16S) | eBioscience | Cat# 11-5773-82 ; RRID: AB_465243 | 200 |
| CD172a Pecy7 (P84) | Biolegend | Cat# 144007; RRID: AB_2563545 | 200 |
| CD172a APC (P84) | BD | Cat# 560106, RRID:AB_1645218 | 200 |
| Streptavidin APC-R700 | BD | Cat# 565144, RRID:AB_2869657 | 800 |
| FLT3 PE (A2F10) | eBioscience | Cat# 12-1351-82, RRID: AB_465859 | 200 |
| ESAM PE (1G8/ESAM) | Biolegend | Cat# 136203; RRID: AB_1953300 | 200 |
| TNFa PE-Cy7 (MP6-XT22) | Biolegend | Cat# 506305; RRID: AB_315426 | 100 |
| IL2 PE-Cy5 (JES6-5H) | Biolegend | Cat# 503824; RRID: AB_2123674 | 100 |
| GranzymeB FITC (QA16A02) | ebioscience | Cat# 11-8898-82; RRID: AB_10733414 | 100 |
| CD4 PerCP-Cy5.5 (GK1.5) | Biolegend | Cat# 100434; RRID: AB_893324 | 400 |
| CD73 PE (eBioTY/11.8) | Biolegend | Cat# 127205; RRID: AB_1089065 | 100 |
| CD90.2 APC (30–H12) | Biolegend | Cat# 105311; RRID: AB_313182 | 100 |

| | | | |
|---|---|---|---|
| CD140a APC (APA5) | Biolegend | Cat# 135908; RRID: AB_2043970 | 100 |
| CCR5 APC (HM-CCR5) | Biolegend | Cat# 107011; RRID: AB_2074528 | 100 |
| CXCR3 (S18001A) PE | Biolegend | Cat# 155903; RRID: AB_2783130 | 100 |
| Anti-mouse CD28, clone 37.51, Ultra-LEAF™ | Biolegend | Cat# 102116; RRID: AB_11147170 | 2000 |
| Anti-mouse CD3e (17A2) | Biolegend | Cat# 100243; RRIB: AB_2563946 | 200 |
| Anti-mouse IL-4 (clone 11B11) Ultra-LEAF™ format | Biolegend | Cat# 504122, RRID: AB_11149679 | 200 |
| InVivoMAb rat IgG2b isotype control, anti-keyhole limpet hemocyanin (clone LTF-2) | BioXcell | Cat# BE0090; RRID: AB_1107780 | See Methods section |
| InVivoMAb anti-mouse agonist PD-1 (B7-H1) | BioXcell | Cat# BE0273; RRID: AB_10949073 Lot: 883023J1 | See Methods section |
| InVivoMAb anti-mouse agonist CTLA4 (CD152, Clone 9H10) | BioXcell | Cat# BE0131; RRID: AB_10950184 Lot: 834323S1 | See Methods section |
| InVivoMab anti-mouse CD4 (GK1.5) | BioXcell | Cat# BE0003-1; RRID: AB_1107636 Lot: 805422A1 | See Methods section |
| InVivoMab anti-mouse CD8 (2.43) | BioXcell | Cat# BE0061; RRID: AB_1125541 Lot: 811522A2 | See Methods section |
| InVivoMab anti-mouse NK1.1 (PK136) | BioXcell | Cat# BE0036; RRID: AB_1107737 Lot: 796521N1 | See Methods section |

| | | | |
|---|---|---|---|
| InVivoMAb anti-mouse CD25 (PC-61.5.3) | BioXcell | Cat# BE0012; RRID:AB_1107619 Lot:795321D1 | See Methods section |
| InVivoMAb anti-mouse CD40 (FGK4.5) | BioXcell | Cat# BE0016-2; RRID:AB_1107647 Lot: 805122F1 | See Methods section |
| CD45 Biotin (30-F11) | Biolegend | Cat# 103103; RRID:AB_312968 | 400 |
| Ly6G Biotin (1A8) | Biolegend | Cat# 127604; RRID: AB_1186105 | 400 |
| CD3 Biotin (145-2C11) | Biolegend | Cat# 100304; RRID: AB_312669 | 400 |
| CD19 Biotin (6D5) | Biolegend | Cat# 115503; RRID: AB_313638 | 400 |
| CD45R/B220 Biotin (RA3-6B2) | Biolegend | Cat# 103203; RRID: AB_312988 | 400 |
| NK1.1 Biotin (PK136) | Biolegend | Cat# 108703; RRID: AB_313390 | 400 |
| SiglecF Biotin (F17007L) | Biolegend | Cat# 155512; RRID: AB_2814066 | 400 |
| TER119 Biotin (TER-119) | Biolegend | Cat# 116203; RRID: AB_313704 | 400 |
| CD11b Biotin | Biolegend | Cat# 101204, RRID:AB_312787 | 400 |
| I-A/I-E Biotin | Biolegend | Cat# 107604, RRID:AB_313319 | 400 |
| **Human antibodies** | | | |
| Human FLT3L Biotin (Polyclonal) | Bio-Techne/R&D systems | Cat# BAF308; RRID:AB_2278494 | 100 |
| NKp46 Biotin (9E2) | Biolegend | Cat#331906, RRID:AB_1027671 | 40 |
| CD3 Biotin (OKT3) | Biolegend | Cat#317319, RRID:AB_10918432 | 400 |
| CD19 Biotin (HIB19) | Biolegend | Cat#302203, RRID:AB_314233 | 400 |
| CD56 Biotin (HCD56) | Biolegend | Cat#318319, RRID:AB_893392 | 40 |
| CD66b Biotin (G10F5) | Biolegend | Cat#305120, RRID:AB_2566608 | 40 |
| CD203c Biotin (REA826) | Miltenyi | Cat#130-112-811, RRID:AB_2656166 | 100 |
| CD20 Biotin (2H7) | Biolegend | Cat#302349, RRID:AB_2565523 | 200 |

| | | | |
|---|---|---|---|
| HLA-DR BUV737 (L243) | BD | Cat#753688, RRID:AB_3687372 | 100 |
| CD163 BUV661 (GHI/61) | BD | Cat#741645, RRID:AB_2871044 | 100 |
| BDCA-2 BUV615 (V24-785) | BD | Cat#751078, RRID:AB_2875114 | 100 |
| CD16 BUV395 (3G8) | BD | Cat#563785 | 100 |
| BTLA BV421 (MIH26) | Biolegend | Cat#344511, RRID:AB_2566507 | |
| CD5 BV605 (L17F12) | Biolegend | Cat#364019, RRID:AB_2565940 | 100 |
| CD123 BV650 (6H6) | Biolegend | Cat#306019, RRID:AB_11218792 | 100 |
| CD141 BV711 (M80) | Biolegend | Cat#344135, RRID:AB_3097423 | 40 |
| CD14 BV786 (M5E2) | BD | Cat#563699 | 100 |
| AXL FITC (108724) | RnD Systems | Cat#18624513 | 100 |
| XCR1 PerCP-Cy5.5 (S15046E) | Biolegend | Cat#372629, RRID:AB_2924564 | 100 |
| CLEC10A PE (H037G3) | Biolegend | Cat#354703, RRID:AB_11219202 | 100 |
| CD88 PE-DAZZLE (S5/1) | Biolegend | Cat#344317, RRID:AB_2750446 | 100 |
| CD45 PE-Cy5 (HI30) | BD | Cat#555484, RRID:AB_395876 | 200 |
| CD1c PE-Cy7 (L161) | Biolegend | Cat#331515, RRID:AB_1953227 | 100 |
| CLEC9A APC (8F9) | Biolegend | Cat#353805, RRID:AB_2565518 | 100 |
| CX3CR1 R718 (2A9-1) | BD | Cat#752200, RRID:AB_2917307 | 100 |
| CD45RA APC-Cy7 (HI100) | Biolegend | Cat#304127, RRID:AB_10708419 | 100 |
| **Chemicals, peptides, and recombinant proteins** | | | |
| Collagenase D | Sigma | Cat# 11088866001 | |
| Collagenase A | Sigma | Cat# 10103586001 | |
| DNase I | Sigma | Cat# 10104159001 | |
| Dispase II | Sigma | Cat# D4693 | |
| HEPES | Gibco | Cat# 11560496 | |

| | | | |
|---|---|---|---|
| Penicillin/Streptomycin | Gibco | Cat# 11528876 | |
| Bovine Serum Albumin Raction V (BSA) | Euromedex | Cat# 04-100-812 | |
| Accucheck counting beads | ThermoFischer | Cat# PCB100 | |
| DMEM Medium, GlutaMAX™ Supplement | Gibco | Cat# 10566024 | |
| RPMI 1640 Medium, GlutaMAX™ Supplement | Gibco | Cat# 61870036 | |
| HBSS, calcium, magnesium | Gibco | Cat# 24020117 | |
| Trypsin-EDTA (0.25%), phenol red | Gibco | Cat# 25200056 | |
| 2-mercaptoéthanol | Gibco | Cat# 11528926 | |
| Fetal Bovine Serum, qualified, Mexico | Gibco | Cat# 12676029 | |
| ACK Lysing Buffer | Gibco | Cat# A1049201 | |
| Ovalbumin peptide SIINFEKL (H-2 Kb) | IVA Lifesciences | Cat# 6-7015-901 | |
| iTAg Tetramer/PE – H-2 Kb OVA (SIINFEKL) | MBL | Cat# TB-5001-1 | 20 |
| Recombinant mouse IL12p70 (carrier-free) | Biolegend | Cat# 577004 | |
| Zombie Yellow™ Fixable | Biolegend | Cat#423104 | 1000 |
| Zombie NIR™ Fixable | Biolegend | Cat#423106 | 1000 |
| Zombie Aqua™ Fixable | Biolegend | Cat#423102 | 1000 |

| | | | |
|---|---|---|---|
| Live/Dead Fixable Blue | ThermoFisher | Cat#L23105 | 1000 |
| 7-AAD | Biolegend | Cat#420404 | 100 |
| Puromycin | Gibco | Cat#A1113803 | |
| DAPI (4',6-Diamidino-2-Phenylindole, Dilactate) | Biolegend | Cat# 422801 | 1000 |
| AMG 487 Antagonist CXCR3 | MedChemExpress | Cat# HY-15319 | |
| Maraviroc Antagonist CCR5 | MedChemExpress | Cat# HY-13004 | |
| Recombinant mouse IL-2 (carrier-free) | Biolegend | Cat# 575402 | |
| Geltrex™ LDEV-Free Reduced Growth Factor Basement Membrane Matrix | Gibco | Cat# A1413201 | |
| PMA (Phorbol 12-myristate 13-acetate) | Sigma-Aldrich | Cat# 16561-29-8 | |
| Ionomycin | Sigma-Aldrich | Cat# 56092-82-1 | |
| Recombinant human Flt3-L | Amgen Inc. | Cat# CDX-301 | |
| Nano-Glo® Fluorofurimazine In Vivo Substrate | Promega | Cat# N4110 | |
| Alanine Aminotransferase (ALT/GPT) Activity Assay Kit | ThermoFisher | Cat# EEA001 | |
| Polybrene | Santa Cruz Biotechnology | Cat# sc-134220 | |
| 4% paraformaldehyde solution | ThermoFisher Scientific | Cat# J19943.K2 | |

| Plasmid and vectors | | |
|---|---|---|
| pMX-IRES-GFP | Origene | NM_001204502.1<br>DOI: 10.1038/s41467-020-15937-y |
| pMX-IRES-Cherry | Origene | NM_000460.2<br>DOI: 10.1038/s41467-020-15937-y |
| pMX-huFLT3L-IRES-GFP | Dr. P. Guermonprez | DOI: 10.1038/s41467-020-15937-y |
| pMX-huFLT3L-IRES-mCherry | This paper | |
| Human CXCL9 Sequence | Cat#: MHS6278-202806039 | Dharmacon horizon |
| Human CCL5 Sequence | Cat#: MHS6278-202757576 | Dharmacon horizon |
| pMX-huCXCL9-IRES-GFP | This paper | |
| pMX-huCCL5-IRES-GFP | This paper | |
| pMX-huCCL5-P2A-huCXCL9-IRES-GFP | This paper | GeneArt Gene Synthesis |
| NanoLuc lentiviral vector | Dr. Laleh Majlessi | N/A |
| **Experimental models: Cell lines** | | |
| B16F10 | | DOI: 10.1016/j.immuni.2020.06.002 |
| B16-huFLT3L-GFP | | DOI: 10.1016/j.immuni.2020.06.002 |
| B16-OVA | This paper | |
| YUMM-OVA | This paper | |
| E0771 | Dr. Stéphanie Hugues | ATCC Number: CRL-3461 |
| TC-1-Luc | Dr. Alexandre Boissonnas | N/A |

| | | |
|---|---|---|
| MC38 | Dr. Philippe Bousso | Cat# SCC172 (Sigma-Aldrich) |
| Mesenchymal stromal cells (MSCs) | Dr. Loredana Saveanu | N/A |
| eMSC-GFP | This paper | |
| eMSC-mCherry | This paper | |
| eMSC-GFP-mCherry | This paper | |
| eMSC-huFLT3L-GFP | This paper | |
| eMSC-huFLT3L-mCherry | This paper | |
| eMSC-huCCL5-GFP | This paper | |
| eMSC-huCXCL9-GFP | This paper | |
| eMSC-huFLT3L-mCherry-huCXCL9-P2A-huCCL5-GFP | This paper | |
| eMSC-NanoLuc | This paper | |
| **Critical commercial assays** | | |
| Human Flt-3 Ligand/FLT3L Quantikine ELISA Kit | R&D | Cat# DFK00 |
| LEGENDplex™ Mouse Proinflammatory Chemokine Panel (13-plex) with V-bottom Plate | Biolegend | Cat# 740451 |
| LEGENDplex™ MU Proinflam. Chemokine Panel 2 (8-plex) | Biolegend | Cat# 741068 |
| Cytofix/cytoperm Kit | BD Bioscience | Cat# 554714 |
| Transcription Factor Staining Buffer Set | ThermoFischer | Cat# 00-5523-00 |

| | | |
|---|---|---|
| EasySep™ Mouse Hematopoietic Progenitor Cell Isolation Kit | Stemcell | Cat# 19856 |
| Human CXCL9 ELISA | R&D Sytstems | Cat# DY392-05 |
| Human CCL5 ELISA | R&D Sytstems | Cat# DY278-05 |
| Corning® Transwell® polycarbonate, ø inserts 6.5 mm, porosity 5 µm (12 inserts in 24-well plate) | Sigma-Aldrich Corning | Cat# 003421 |
| **Experimental models: Organisms/strains** | | |
| mouse: C57BL/6J | Janvier Labs | RRID:IMSR_JAX:000664 |
| mouse: CD45.1 (CByJ.SJL(B6) *Ptprc*$^{a}$/J) | Janvier Labs | RRID:IMSR_JAX:006584 |
| mouse: OT-1 Rag2-/- CD45.1 | Dr. Sebastian Amigorena | DOI: 10.1038/s41467-022-31504-z |
| mouse: Rosa-DTA (B6.129P2 *Gt(ROSA)26Sor*$^{tm1(DTA)Lky}$/J) | Dr. Marc Dalod | RRID:IMSR_JAX:009669 |
| mouse: ROSA26-LSL-RFP (B6.Cg *Gt(ROSA)26Sor*$^{tm1Hjf}$/J) | Dr. Marc Dalod | RRID:IMSR_JAX:038164 |
| mouse: *Xcr1-Cre* (B6 *Xcr1*$^{tm1Ciphe}$) | Dr. Marc Dalod | DOI: 10.3389/fimmu.2018.02805 |
| Mouse: ROSA26-LSL-tdTomato (B6.Cg *Gt(ROSA)26Sor*$^{tm14(CAG-tdTomato)Hze}$/J) | Dr. Tessa Bergsbaken | RRID:IMSR JAX:007914 |

| | | |
|---|---|---|
| mouse: *Irf8*-GFP (B6.Cg *Irf8*$^{tm2.1Hm}$/J) | The Jackson Laboratory | RRID: IMSR_JAX:027084 |
| mouse: BALB/c *Rag2*$^{-/-}$ *IL2rγ*$^{-/-}$ *Sirpα*$^{NOD}$ (BRGS), BALB/c *Rag2*$^{-/-}$ *IL2rγ*$^{-/-}$ *Sirpα*$^{NOD}$ *Flt3*$^{+/+}$ | Human Disease Model Core facility - Pasteur Institute | DOI : 10.1002/eji.201848001 |
| mouse: BALB/c *Rag2*$^{-/-}$ *IL2rγ*$^{-/-}$ *Sirpα*$^{NOD}$ (BRGS), BALB/c *Rag2*$^{-/-}$ *IL2rγ*$^{-/-}$ *Sirpα*$^{NOD}$ *Flt3*$^{-/-}$ | Human Disease Model Core facility - Pasteur Institute | DOI : 10.1002/eji.201848001 |
| **Software and algorithms** | | |
| BD FACSDiva | BD | |
| FlowJo v10.10 | BD | https://www.flowjo.com |
| GraphPad Prism 10 | GraphPad Software | https://www.graphpad.com |
| RStudio 4.4.2 | The R Foundation | https://www.r-project.org |
| Morpheus | Broad institute | https://software.broadinstitute.org/morpheus/ |
| LEGENDplex™ Data Analysis Software | Biolegend | https://legendplex.qognit.com/ |
| NIS-Elements AR 6. 10 | Nikon | |
| Image J v1. 54f | Schneider et al., 2012 | https://imagej.nih.gov/ij/ |
| QuPath 0.4.4 | Qupath software | https://qupath.github.io/ |
| ID7000 Software 2.2.1.17271 | Sony | |
| Living Image 4.8.2 Software | Caliper Life Sciences | |

| | |
|---|---|
| Gen5 3.12 software | Agilent BioTek |
